# Targeting Reactive Aldehyde Detoxification by Aldehyde Dehydrogenase 2 (ALDH2) as a Treatment Strategy for Endometriosis

**DOI:** 10.1101/2020.01.30.927376

**Authors:** Stacy L. McAllister, Pritam Sinharoy, Megana Vasu

## Abstract

Endometriosis affects ∼176 million women worldwide, yet on average, women experience pain ∼10 years from symptom onset before being properly diagnosed. Standard treatments (drugs or surgery) often fail to provide long-term pain relief. Elevated levels of reactive aldehydes such as 4-hydroxynonenal (4-HNE) have been implicated in the peritoneal fluid of women with endometriosis and upon accumulation, reactive aldehydes can form protein-adducts and/or generate pain. A key enzyme in detoxifying reactive aldehydes to less reactive forms, is the mitochondrial enzyme aldehyde dehydrogenase-2 (ALDH2). Here, we tested the hypothesis that aberrant reactive aldehyde detoxification by ALDH2, underlies endometriosis and its associated pain. We determined, in the eutopic and ectopic endometrium of women with severe (stage IV) peritoneal endometriosis, that ALDH2 enzyme activity was decreased, which was associated with decreased ALDH2 expression and increased 4-HNE adduct formation compared to the eutopic endometrium of controls in the proliferative phase. Using a rodent model of endometriosis and an ALDH2*2 knock-in mouse with decreased ALDH2 activity, we determined that increasing ALDH2 activity with the enzyme activator Alda-1 could prevent endometriosis lesion development as well as alleviate pain-associated behaviors in proestrus. Overall, our findings suggest that targeting the ALDH2 enzyme in endometriosis may lead to better treatment strategies and in the proliferative phase, that increased 4-HNE adduct formation within the endometrium may serve as a less invasive diagnostic biomarker to reduce years of suffering in women.

**One Sentence Summary:** ALDH2 activity influences endometriosis and its associated pain.

## Introduction

Endometriosis is an estrogen dependent inflammatory condition defined by the growth of endometrial tissue in extrauterine locations (variously called lesions, cysts, ectopic growths, implants). The condition affects ∼176 million women worldwide, yet little progress has been made over the past 20 years relative to screening, detection, prognosis, and treatment (1). The most common symptom of endometriosis is pain and 70-90% of women of reproductive age with chronic pelvic pain (CPP), have endometriosis (2). Painful symptoms include debilitating pelvic/abdominal pain, dyspareunia (vaginal hyperalgesia, pain during intercourse), severe dysmenorrhea (pain on menstruation), dyschezia (pain on defecation), and dysuria (pain with urination). Women with the condition also suffer from co-occurring painful conditions including interstitial cystitis/painful bladder syndrome, irritable bowel syndrome, vulvodynia, fibromyalgia, and up to 50% of these women also suffer from infertility (3, 4). A major clinical problem is that painful symptoms associated with endometriosis poorly correlate with disease extent and on average, women experience pain ∼10 years before being properly diagnosed (5-7). The gold standard for endometriosis diagnosis is laparoscopic visualization of the lesions preferably with histological confirmation, which is invasive and expensive. Available treatments for endometriosis include drugs and/or surgery, which tend to be ineffective over the long-term and can produce unwanted side effects premature bone loss, vaginal dryness, and contraception. Thus, there is a need for more effective pain therapeutics and less invasive diagnostic strategies to reduce years of suffering in women with endometriosis.

How endometriosis occurs is not fully understood and is considered an enigma. The leading hypothesis, Sampson’s hypothesis, suggests retrograde menstruation underlies the disease, in which the endometrial lining (endometrium) of the uterus travels retrogradely through the fallopian tubes and implants, primarily in the peritoneal cavity (8). However, ∼90% of women experience retrograde menstruation but only ∼10% have endometriosis, suggesting differences within the endometrium, possibly genetic, underlie the ability of the ectopic lesion to successfully implant, progress, and avoid immune system clearance (9). One factor known to be involved in the pathogenesis, progression, and establishment of endometriosis is oxidative stress (10, 11). Under oxidative stress, excess reactive oxygen species (ROS) are produced and as a secondary bi-product of lipid peroxidation, reactive aldehydes including 4-hydroxynonenal (4-HNE), are generated (12). In the peritoneal fluid of women with endometriosis, elevated reactive aldehyde levels have been implicated and through accumulation, reactive aldehydes can form protein adducts and/or generate pain (13-19). A critical enzyme in detoxifying reactive aldehydes such as 4-HNE to unreactive forms is the mitochondrial enzyme ALDH2. Here, we tested the hypothesis that aberrant reactive aldehyde detoxification by ALDH2, underlies the painful condition of endometriosis.

## Results

### Women with endometriosis have decreased ALDH2 activity

To determine how ALDH2 activity is regulated in the endometrium of women with endometriosis, we analyzed and compared endometrial biopsies from women with severe (stage IV) peritoneal endometriosis (eutopic and patient-matched ectopic endometrium) and without endometriosis (eutopic) collected in the proliferative phase (see Table 1 for patient and biopsy characteristics). In the eutopic and ectopic endometrium of women with endometriosis, we determined that ALDH2 enzyme activity was decreased (*P* < 0.005 and *P* < 0.0005, respectively) which was associated with decreased (*P* < 0.0005 and *P* < 0.0005, respectively) ALDH2 protein expression (Fig. 1A). To determine if decreased reactive aldehyde detoxification by ALDH2 was associated with increased protein-adduct formation, we assessed the same endometrial biopsies for 4-HNE adducts. We determined, in women with endometriosis, that 4-HNE adduct formation in the eutopic and ectopic endometrium was increased (*P* < 0.05 and *P* < 0.05, respectively), compared to the eutopic endometrium of women without endometriosis (Fig. 1B-C). No significant differences were found between the eutopic and ectopic endometrium, relative to ALDH2 activity, expression, or 4-HNE adduct formation in women without endometriosis (Fig. 1A-C). Together, these findings suggest that decreased ALDH2 activity and expression may underly endometriosis pathophysiology and that increased 4-HNE adduct formation in the endometrium may occur as a result of decreased reactive aldehyde detoxification by ALDH2, supporting our hypothesis.

**TABLE 1.** Sample Characteristics.

| Sample ID | Cycle phase | Age (yr) | Race | Weight (kg) | BMI (kg/m<sup>2</sup>) | Medications/Other |
| --- | --- | --- | --- | --- | --- | --- |
| **Severe stage (IV) peritoneal endometriosis** |  |  |  |  |  |  |
| SE178 | PE | 47 | Caucasian | 84.55 | 46.65 | Lexapro, Wellbutrin |
| SE182 | PE | 39 | Caucasian | 68.18 | 28.40 | Zyrtec, Prozac |
| SE183 | PE | 29 | Caucasian | 61.36 | 22.51 | None |
| EN141 | PE | 43 | Caucasian | 85.73 | 33.48 | Albuterol, Nasonex, Tylenol |
| EE185 | PE | 31 | Caucasian | 50.91 | 21.21 | Ritalin |
| **No endometriosis** |  |  |  |  |  |  |
| EN142 | PE | 49 | Caucasian | 79.38 | 32.01 | Albuterol, Zantac, Claritin |
| EN146 | PE | 38 | Caucasian | 58.97 | 24.56 | Allegra, Vitamins |
| EN150 | PE | 36 | Caucasian | 56.70 | 20.80 | Levothyroxine, Advil, Vitamins |
| EN152 | PE | 41 | Caucasian | 77.56 | 27.60 | Lexapro |
| SN 134 | PE | 43 | Caucasian | 123.64 | 46.79 | Lovenox, Coumadin, Norco, Colace, Iron |

All subjects were nonsmokers (except SE178 was past smoker) and were not on steroid hormone medications within 3 months prior to tissue sampling. PE, Proliferative phase of the menstrual cycle, determined by histologic evaluation according to Noyes criteria. Peritoneal endometriosis, defined as biopsy-proven serosal implant. Severe endometriosis (Stage IV disease), defined in accordance with the Revised American Fertility Society (rAFS) classification system.

**Fig 1.**
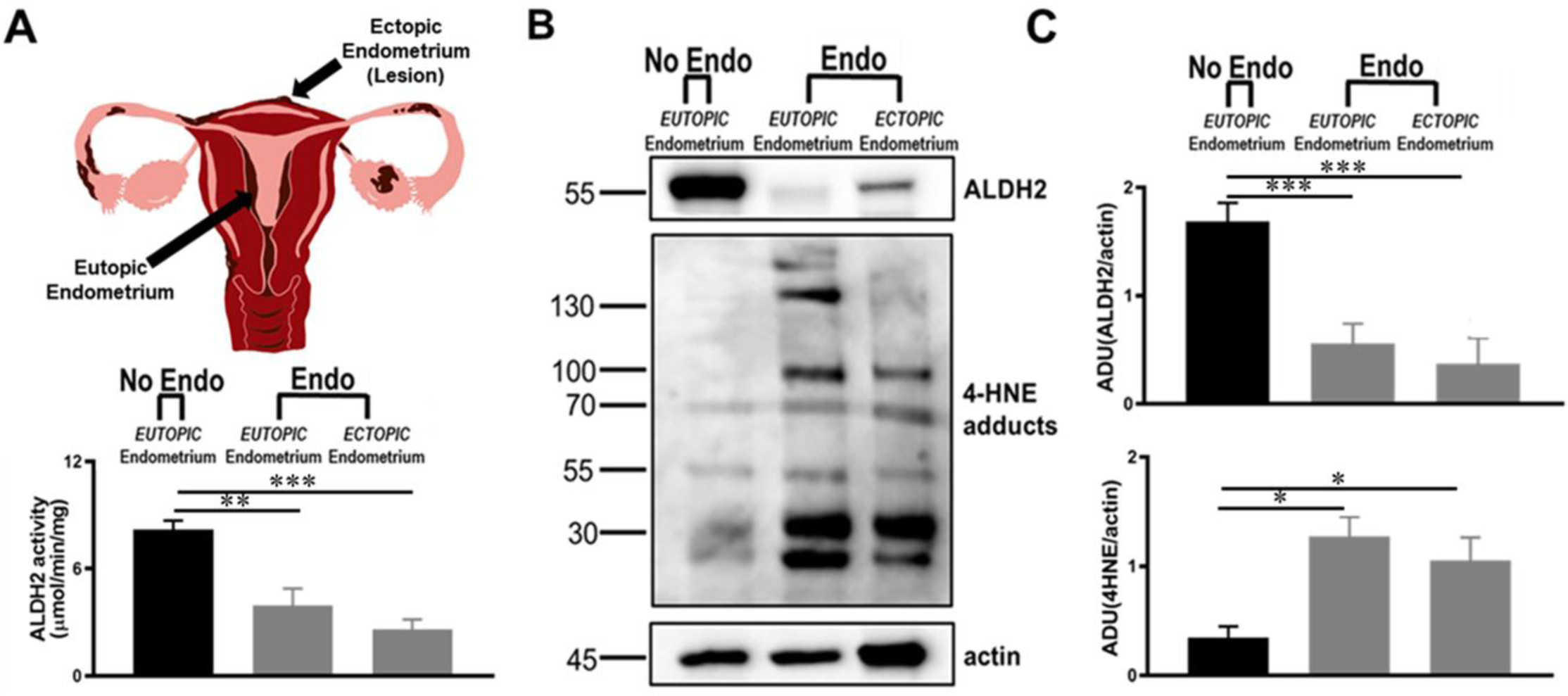
Molecular characterization of the endometrium of women without and with endometriosis. In the proliferative phase, ALDH2 activity of the endometrium was analyzed in women without endometriosis (No Endo: eutopic) and in women with severe (IV) peritoneal endometriosis (Endo: eutopic and ectopic endometrium, patient matched) aud expressed as μmol/min/mg. The reduction of NAD+ to NADH at λ340 nm was measured using a spectrophotometer and 25mM acetaldehyde as a substrate. Data presented is absorbance measured during the first 2 minutes after acetaldehyde substrate was added. (A). **B-C.** Representative western blot of ALDH2 and 4-HNE adduct formation in the endometrium of women without endometriosis (eutopic) and with endometriosis (eutopic and ectopic lesion) (B). Western blot analysis of ALDH2 expression (**top**) and 4-HNE adduct formation (**bottom**) relative to actin as loading control (C). All data are expressed as mean ±SEM, n=5 biological replicates/group. Assessed by one-way ANOVA followed by Tukey post hoc test. *p<0.05, **p<0.005. ***p<0.0005

### Female naïve ALDH2*2 homozygote knock-in mice have decreased ALDH2 activity

To determine if an ALDH2*2 knock-in mouse with decreased ALDH2 activity can be used as a unique tool for studying endometriosis, we biochemically characterized ALDH2 activity, ALDH2 protein expression, and 4-HNE adduct formation. Using an enzyme activity assay, we determined that relative to wild type mice, ALDH2*2 mice had ∼70% and 63% reduced liver (control) and uterus ALDH2 activity, respectively (Fig. 2A). Western blot revealed that, relative to wild type mice, ALDH2*2 mice had ∼60% and 62% reduced ALDH2 protein expression in the liver and uterus, respectively (Fig. 2B and C). Under physiological conditions, no significant differences in 4-HNE adduct formation were found between naïve wild type and naïve ALDH2*2 mice liver and/or uterus, suggesting similar redox states in the absence of disease pathology (Fig. 2B and C). Together, these findings suggest that the ALDH2*2 mouse, with decreased ALDH2 activity and expression relative to wild type, and similar basal levels of 4-HNE adduct formation, is a unique tool to begin to elucidate the role of reactive aldehyde detoxification by ALDH2 in endometriosis.

**Fig 2.**
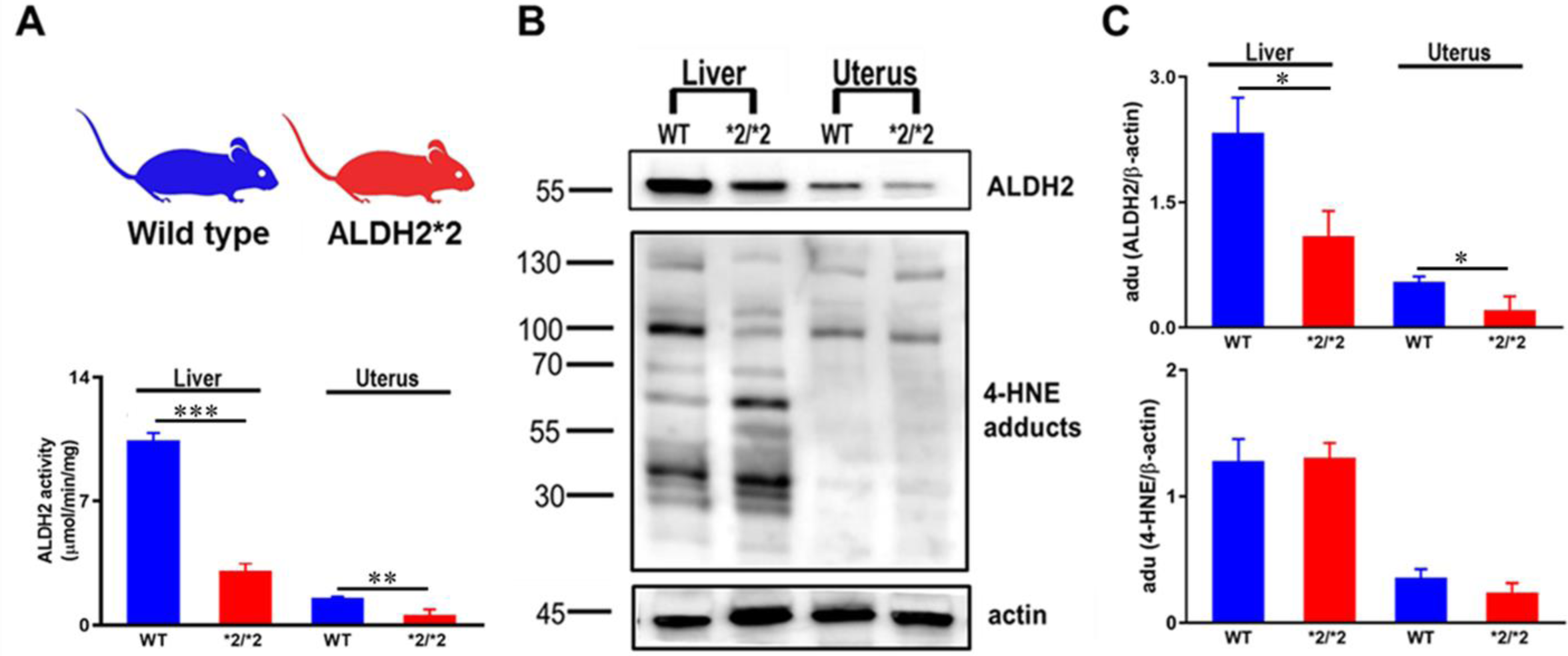
Molecular characterization of naïve female wild type and homozygotic ALDH2*2 knock-in mice. Wild type C57BL/6 mice (blue) and ALDH2*2 knock-in mice (red) with a C57BL/6 background. The ALDH2*2 knock-in mouse has a single amino acid substitution in which adenine is substituted for guanine at the first base pair of codon 487. The result is an amino acid change from glutamic acid (Glu, E) to lysine (Lys, K) that is equivalent to the E487K substitution in the human ALDH2*2 variant. In proestrus, mouse liver (control) and uterus ALDH2 enzyme activity was spectrophotometrically determined by measuring the reduction of NAD+ to NADH at λ340 nm using 25mM acetaldehyde as a substrate. Data presented is absorbance measured during the first 2 minutes after acetaldehyde substrate was added. Activity is expressed as μmol/min/mg (A). **B-C.** Representative western blot of ALDH2 protein expression and 4-HNE adduct formation in liver and uterus of wild type and ALDH2*2 mice (B). Western blot analysis of liver and uterus ALDH2 protein expression (**top**) and 4-HNE adduct formation (**bottom**) relative to actin loading control. All data are expressed as mean ±SEM, n=5 biological replicates/group. Assessed by two-tailed Student’s *t*-test test, blue and red bars indicate wild type and ALDH2*2 mice, respectively. *p<0.05, **p<0.0005, ***p<0.0001

### Decreased ALDH2 activity accelerates endometriosis development in a rodent model

To determine if decreased ALDH2 enzyme activity contributes to lesion development, endometriosis was induced in wild type and ALDH2*2 mice using a validated rodent model that produces signs (fluid filled, vascularized, innervated lesions) and painful symptoms similar to that of women with endometriosis (Fig. 3A) (21-25). To establish and compare developmental time courses, mice were sacrificed, lesions measured, and average lesion area determined for both wild type and ALDH2*2 mice at one of four time points post-endometriosis induction: day 1, 3, 14, or 28 (Fig. 3B). By day 3, ALDH2*2 mice developed a larger (*P* < 0.05) lesion area compared to the lesion area at day 1 in ALDH2*2 mice (Fig. 3C). Whereas, in wild type mice, a larger (*P* < 0.0001) lesion area was not observed until day 14, relative to wild type lesion area day 1. At day 28, lesion area in both wild type and ALDH2*2 mice was larger (*P* < 0.0001 and *P* < 0.0001, respectively) than their respective day 1 lesion areas, but not their respective day 14 lesions when lesion development stabilizes and endometriosis is established. Over the developmental time course, comparing wild type and ALDH2*2 mice, at day 3, ALDH2*2 mice developed a larger (*P* < 0.05) lesion area compared to wild type mice, but at day 14 and day 28, no significant differences in lesion area were found between ALDH2*2 and wild type mice (Fig. 3C). These findings suggest that the decreased ALDH2 activity of the ALDH2*2 mice, accelerates early lesion development in endometriosis, relative to wild type mice.

**Fig 3.**
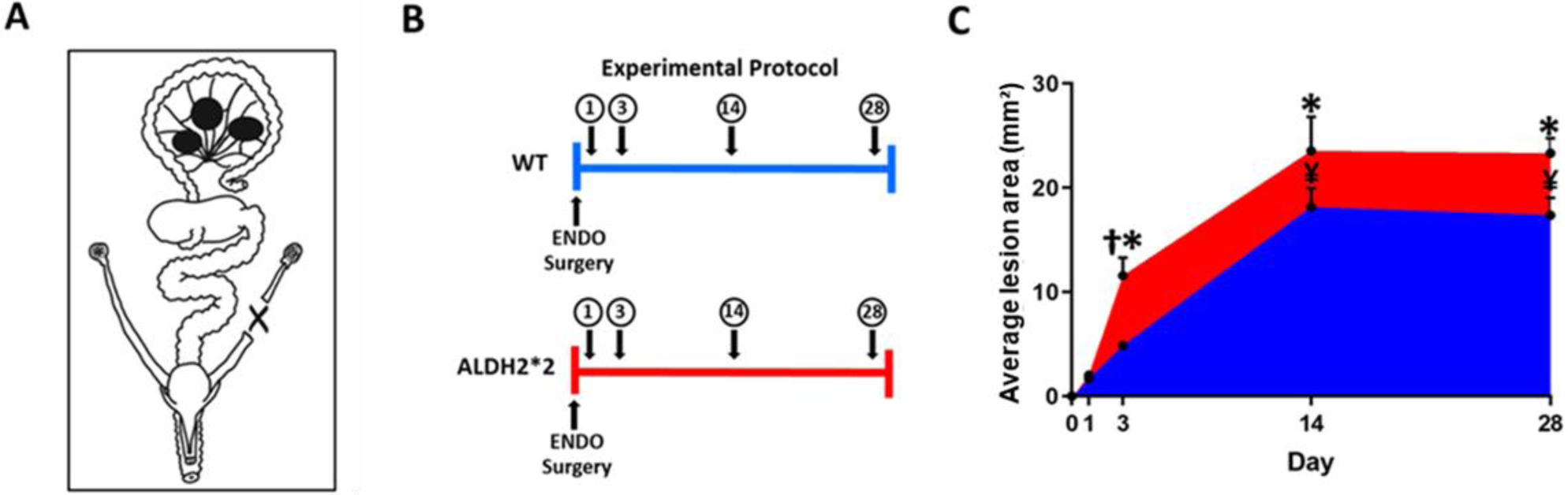
The influence of decreased ALDH2 activity on endometriosis development in a rodent model. Three equal pieces of eutopic uterus with endometrium are sutured onto alternate cascading mesenteric arteries. These auto-transplants form ectopic lesions and symptoms similar to women with endometriosis (A). **B-C.** Experimental protocol: endometriosis surgery was performed in wild type (n=24) and ALDH2*2 (n=24) female mice and then at day 1, 3, 14, or 28 mice sacrificed and lesions measured (n=6/group/genotype) (B). The average lesion area was determined for wild type and ALDH2*2 mice at each time point in the developmental time course (C). All data are expressed as mean ±SEM. Assessed by two-way ANOVA followed by Tukey post hoc test, blue and red bars indicate wild type and ALDH2*2 mice, respectively. **†**p<0.05 *vs*. wild type day 3, *p<0.0001 *vs*. ALDH2*2 day 1, ¥p<0.0001 *vs*. wild type day 1

### Decreased ALDH2 enzyme activity exacerbates endometriosis pain-associated behavior in a rodent model

To determine if decreased ALDH2 enzyme activity contributes to endometriosis pain-associated behaviors, behavioral assessments were made in proestrus in wild type and ALDH2*2 mice for 2 weeks before endometriosis was induced and then during the 2 week time period that endometriosis becomes established (Fig. 4A). At baseline, no significant differences were observed between wild type and ALDH2*2 mice relative to abdominal directed licking (events in 5 min period), paw withdrawal threshold, or thermal latency (Fig. 4B-D). Post-endometriosis, compared to baseline, both wild type and ALDH2*2 mice had increased (*P* < 0.0005 and *P* < 0.0001) abdominal directed licking, decreased (*P* < 0.0001 and *P* < 0.0001) paw withdrawal thresholds, and decreased (*P* < 0.0001 and *P* < 0.0001) thermal latencies, relative to their respective baselines. Post-endometriosis, ALDH2*2 mice compared to wild type mice, had increased (*P* < 0.05) abdominal licking suggesting that the decreased ALDH2 activity exacerbated local abdominal discomfort (Fig. 4B). To assess for differences in locomotor activity, an automated Opto-Varimex activity monitor recorded the total, ambulatory, and vertical counts in a 5 min period (Fig. 4E and fig. S1A and B). No significant differences in locomotor activity were observed between wild type and ALDH2*2 mice at baseline, or within groups post-endometriosis relative to baseline, or between groups post-endometriosis. To assess for differences in exploratory behavior, a modified home cage open field setting with a cardboard tunnel was used. The total time spent in the tunnel (s), number of tunnel entries, and the number of times the mouse climbed on top of the tunnel was recorded in a 5 min period (Fig. 4F and fig. S1C and D). At baseline, no significant differences were observed between wild type and ALDH2*2 mice relative to total time in tunnel (s) or number of times on top of the tunnel, but ALDH2*2 mice had a decreased (*P* < 0.05) number of tunnel entries, relative to wild type mice (fig. S1C). Post-endometriosis, in ALDH2*2 mice, total time in tunnel (s), compared to baseline, was significantly decreased (*P* < 0.005; Fig. 4F). No other significant differences were found between or within groups relative to exploratory behavior. Overall, these findings suggest that the decreased ALDH2 activity of the ALDH2*2 mouse exacerbates endometriosis abdominal pain-associated behavior.

**Fig 4.**
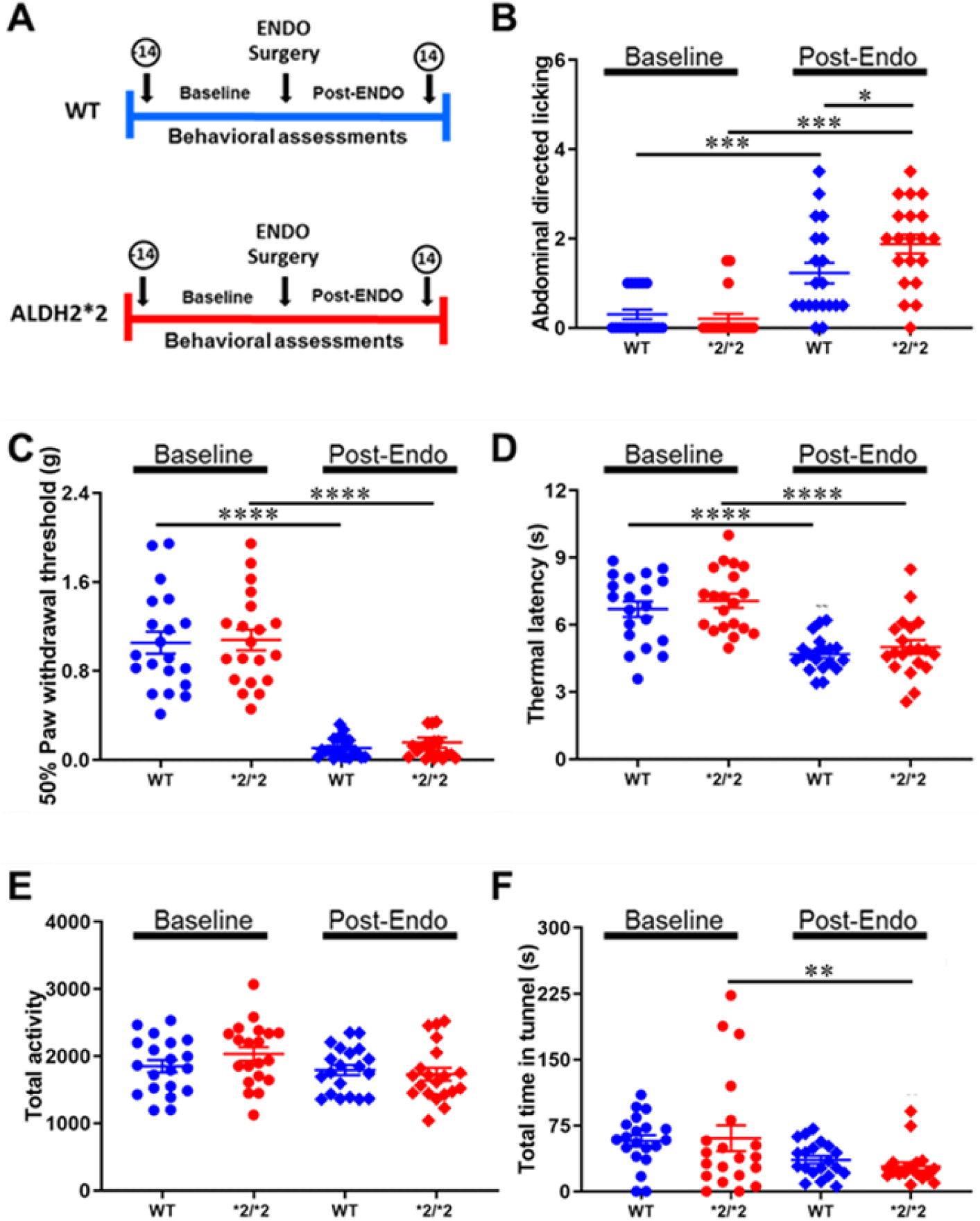
The influence of decreased ALDH2 activity on endometriosis pain-associated behaviors in a rodent model. **A-F.** Experimental Protocol: Behavioral parameters were assessed for 2 weeks before and after endometriosis surgery in wild type (n=20) and ALDH2*2 mice (n=20) in proestrus (∼day 5 and day 13 of both time periods) (A). As a measure of abdominal discomfort or primary pain, the number of times the mouse licked, groomed, or barbered the abdominal region was recorded (B). As an indicator of secondary referred pain, changes in nociception were assessed. Hind paw mechanical withdrawal threshold (C) and thermal latency (D) were assessed by von Frey hairs and Hargreaves method, respectively. Locomotor activity and exploratory behavior were measured by the total activity (E) and total time spent in tunnel (F). Abdominal licking, locomotor, and exploratory behavior were assessed in 5 minutes sessions. Abdominal licking and exploratory behavior were assessed in a modified home cage open field setting. All data are expressed as mean ±SEM. Assessed by 2-way ANOVA with Tukey’s post hoc test, blue and red data points indicate wild type mice, red data points indicate ALDH2*2/*2 mice, circles baseline assessments, diamonds post-endometriosis assessments. *p<0.05, **p<0.005, ***p<0.0005, ****p<0.0001.

### Increasing ALDH2 activity prevents endometriosis development in a rodent model

To determine if increasing ALDH2 enzyme activity could prevent lesion development, we incorporated with our rodent endometriosis model, the small molecule Alda-1[N-(1,3-benzodioxo-5-ylmethyl)-2,6-dichlorbenzamide] that selectively increases ALDH2 activity by correction of the ALDH2*2 mutant structural deficit (20). Beginning the day of endometriosis surgery, in wild type and ALDH2*2 mice, Alzet osmotic pumps were implanted for continuous delivery of Alda-1 (5mg/kg) or DMSO PEG50/50% (control). Mice were then sacrificed, lesions measured, and average area determined for both wild type and ALDH2*2 mice at one of two developmental time points: day 3 or 28 (Fig. 5A). By day 3, ALDH2*2 mice treated with Alda-1 had smaller (*P* < 0.0005) lesion areas compared to DMSO treated ALDH2*2 mice (Fig. 5B). By day 28, both wild type and ALDH2*2 mice treated with Alda-1 had smaller (*P* < 0.005 and *P* < 0.05, respectively) lesion areas compared to respective DSMO treated groups (Fig. 5C). Comparing wild type and ALDH2*2 mice, treated with DMSO, at day 3, ALDH2*2 mice lesion area was larger (*P* < 0.05) than that of wild type mice (Fig. 6A) but by day 28, no significant differences were found between DMSO treated wild type and ALDH2*2 mice relative to lesion areas. Comparing wild type and ALDH2*2 mice, treated with Alda-1, no significant differences in lesion area were found at day 3 or day 28 (Fig. 5C and D). Overall, these findings suggest that Alda-1 prevents the accelerated lesion development seen at day 3 in ALDH2*2 mice, relative to wild type mice (Fig. 3C), and that by day 28, Alda-1 prevents lesion development in both wild type and ALDH2*2 mice, supporting the involvement of ALDH2 activity in endometriosis development.

**Fig 5.**
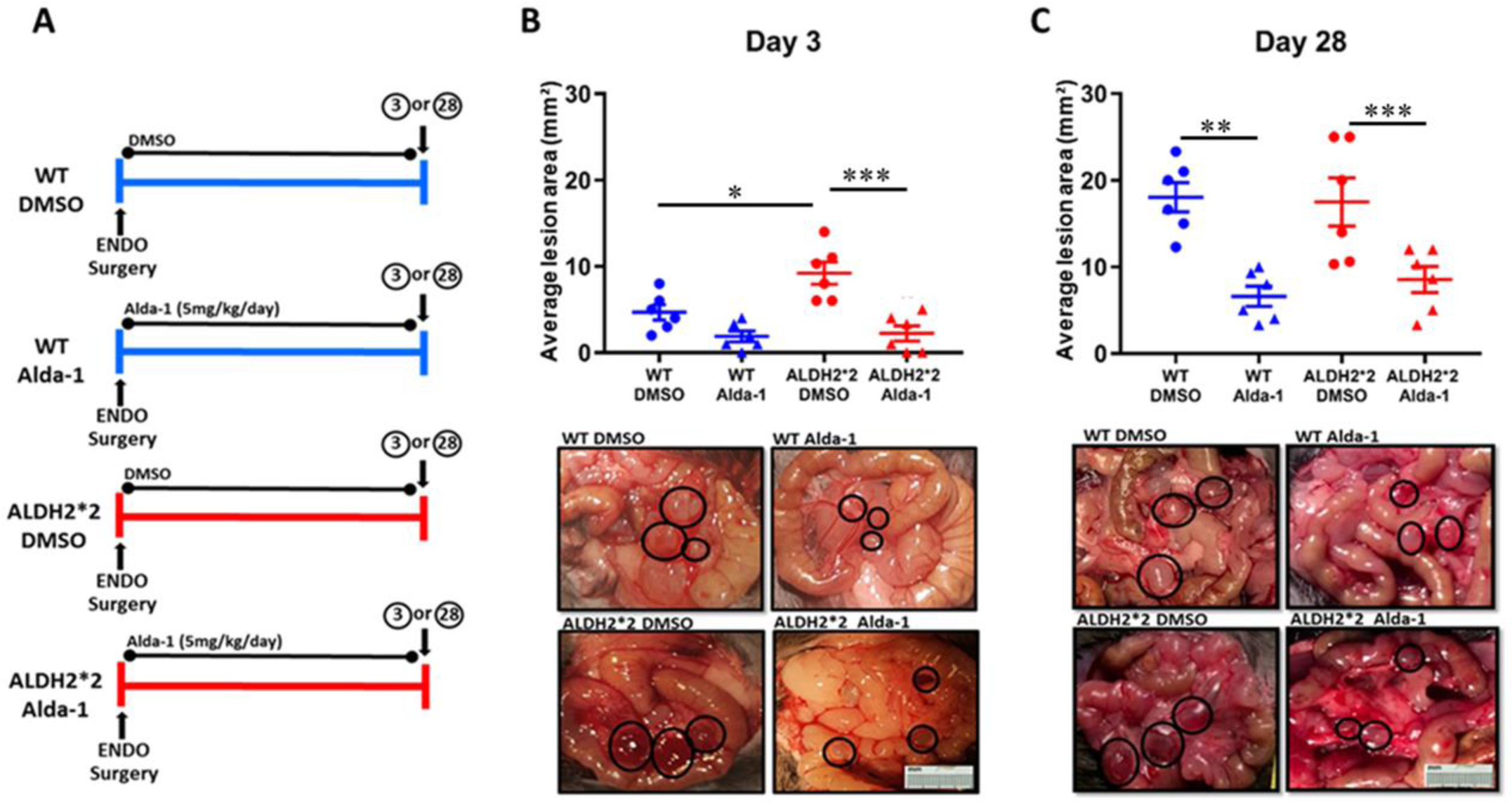
The influence of increased ALDH2 activity on endometriosis development in a rodent model. A-C. Experimental protocol: endometriosis surgery was performed in female wild type (n=24) and ALDH2*2 mice (n=24). Mice were treated for 3 days or 28 days with Alda-1 (5mg/kg/day or as a control 50-50 DMSO-PEG) via Alzet pump delivery beginning the day of surgery (n=6/group/genotype/treatment) (A); Average lesion area and representative lesions in wild type and ALDH2*2 mice treated with Alda-1 or DMSO control for 3 days (B) or 28 days (C). All data are expressed as mean ±SEM. Assessed by one-way ANOVA followed by Tukey post hoc test, blue data points and bars indicate wild type mice, red data points and bars indicate ALDH2*2 mice, circles DMSO treated, triangles Alda-1 treated. *p<0.05, **p<0.005, ***p<0.0005.

**Fig 6.**
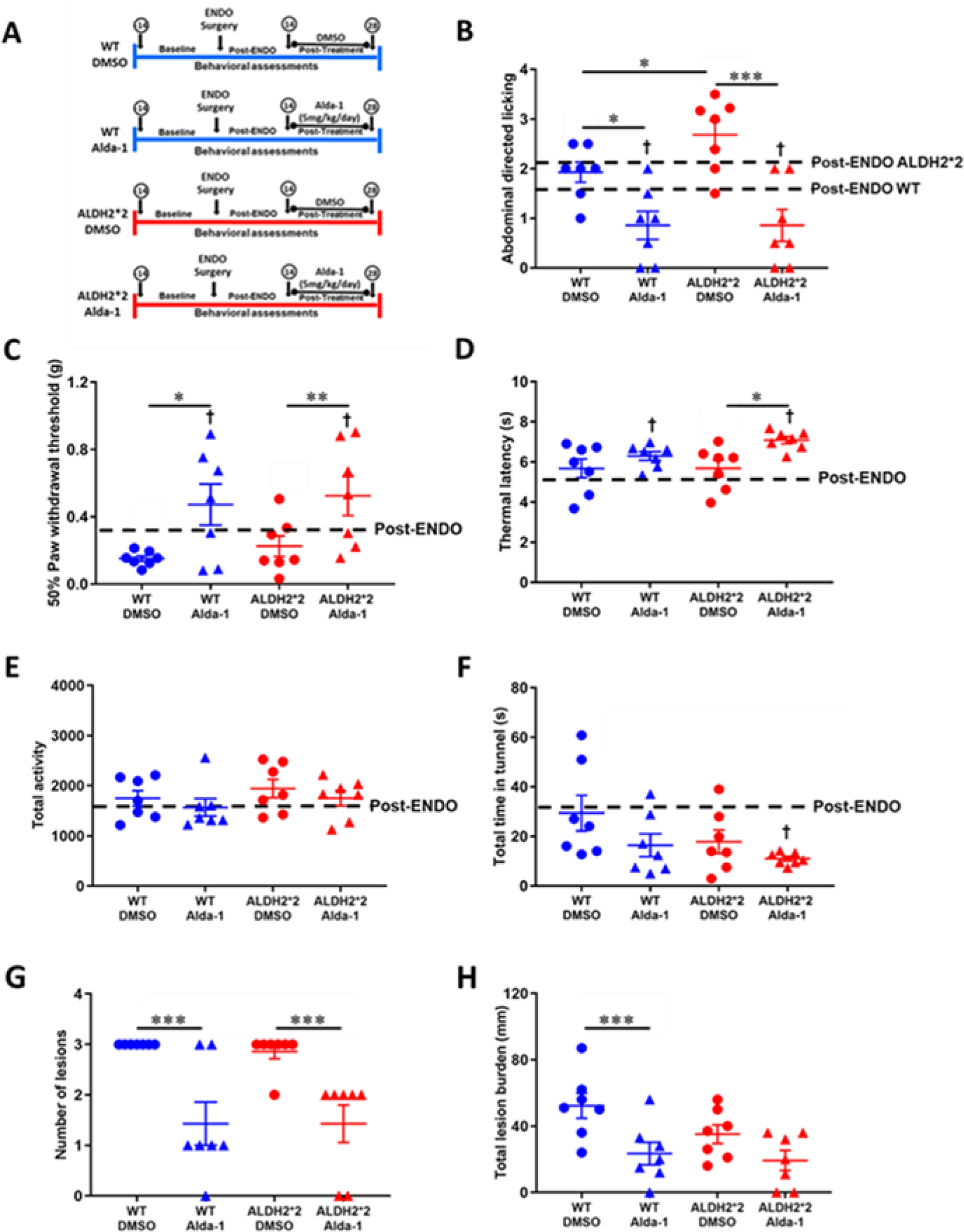
The influence of increased ALDH2 activity on endometriosis pain-associated behaviors in a rodent model. **A-F.** Experimental Protocol: Endometriosis surgery was performed in wild type (n=14) and ALDH2*2 mice (n=14). At day 14 post-endometriosis, Alda-1 treatment (5mg/kg/day or 50-50 DMSO-PEG control) began and continued for 2 weeks via Alzet osmotic pump (n=7/group/genotype) (A). Presented behavioral parameters were assessed in proestrus 2 weeks post-endometriosis surgery and 2 weeks post-treatment (∼day 5 and day 13 of both time periods). Abdominal licking (B), paw withdrawal threshold (C), thermal latency (D), total activity (E), and total time in tunnel (F) were assessed. At the time of sacrifice lesions were assessed to compare number of lesions (A), and total lesion burden (B) between groups. All data are expressed as mean ±SEM. Behavioral data assessed by two-way ANOVA followed by Tukey post hoc test. Lesion data assessed by one-way ANOVA, followed by Tukey post hoc test. Blue data points indicate wild type mice, red data points indicate ALDH2*2/*2 mice, circles DMSO treated, triangles Alda-1 treated. †p<0.05 *vs* post-endometriosis, *p<0.05, **p<0.005, ***p<0.0001

### Increasing ALDH2 activity alleviates endometriosis pain-associated behaviors in a rodent model

To determine if increasing ALDH2 enzyme activity could alleviate pain-associated behaviors once endometriosis is established, Alda-1 (or DMSO PEG50/50% control) treatment was delivered via Alzet osmotic pump beginning 2 weeks post-endometriosis and continued for 2 weeks (Fig. 6A). Behavioral assessments were made before and after endometriosis was induced and then again post-treatment. Compared to post-endometriosis assessments, Alda-1 treated wild type and ALDH2*2 endometriosis mice had decreased (*P* < 0.05 and *P* < 0.05, respectively) abdominal directed licking, increased (*P* < 0.05 and *P* < 0.005, respectively) mechanical paw withdrawal threshold, and increased (*P* < 0.005 and *P* < 0.05, respectively) thermal latency (Fig.6B-D). In DMSO treated wild type and ALDH2*2 endometriosis mice, no significant differences were found in abdominal directed licking, paw withdrawal threshold, or thermal latency compared to their respective post-endometriosis assessments (Fig.6B-D). Alda-1 treated wild type and ALDH2*2 endometriosis mice, compared to their respective DSMO groups, had decreased (*P* < 0.05 and *P* < 0.0001, respectively) abdominal directed licking and an increased (*P* < 0.005 and *P* < 0.005, respectively) mechanical paw withdrawal threshold (Fig. 6B and C). Relative to thermal latency, no significant differences were found between wild type endometriosis mice treated with DMSO or Alda-1; however, Alda-1 treated ALDH2*2 endometriosis mice, had an increased (*P* < 0.05) thermal latency compared to their respective DMSO group (Fig. 6D).

To determine if Alda-1’s alleviation of endometriosis pain-associated behaviors was related to locomotor activity, post-treatment total, ambulatory, and vertical counts were compared to post-endometriosis assessments. No significant differences were found post-treatment in wild type or ALDH2*2 mice treated with Alda-1 or DMSO compared to their respective total, ambulatory, and vertical post-endometriosis counts (Fig. 6E, fig. S2A and B) suggesting Alda-1’s alleviation of pain was independent of locomotor activity. Further, no significant differences were found between Alda-1 treated wild type and ALDH2*2 endometriosis mice, compared to their respective DMSO groups, in locomotor activity.

To determine if Alda-1 treatment influenced exploratory behavior, the total time spent in the tunnel (s), number of tunnel entries, and the number of times the mouse climbed on top of the tunnel, assessments post-treatment were compared to post-endometriosis (Fig. 6F, fig. S2C and D). Alda-1 treated ALDH2*2 mice, compare to post-endometriosis, had a decrease in total time in tunnel and number of tunnel entries (*P* < 0.05 and *P* < 0.05, respectively), suggesting Alda-1 pain alleviation in ALDH2*2 mice may be associated with decreased exploratory behavior (Fig. 6F, fig. S2C). No other significant differences were found in number of tunnel entries, total time in tunnel, or number of times on top of the tunnel in Alda-1 or DMSO treated wild type and ALDH2*2 endometriosis mice, relative to respective post-endometriosis assessments (Fig. 6F, fig. S2C and D). Further, no significant differences in locomotor activity were found between Alda-1 treated wild type and ALDH2*2 endometriosis mice, compared to their respective DMSO groups. Overall, these findings suggest that increasing ALDH2 activity with Alda-1, can alleviate endometriosis pain-associated behaviors in both wild type and ALDH2*2 mice without influencing locomotor activity but in ALDH2*2 mice, exploratory behavior may be influenced.

To determine if Alda-1’s alleviation of endometriosis pain-associated behaviors was associated with lesion parameters, the average area, number of lesions, and total lesion burden was analyzed and compared post-treatment. In wild type and ALDH2*2 mice treated with Alda-1, no significant differences were found in average lesion area, relative to respective DMSO treated mice (fig. S2E). However, in both Alda-1 treated wild type and ALDH2*2 mice, the number of lesions were decreased (*P* < 0.0001 and *P* < 0.0001, respectively) compared to respective DMSO treated mice (Fig. 6G). To account for the reduced lesion number, the total lesion burden or total amount of ectopic growth for each group was analyzed and compared. In Alda-1 treated wild type mice, total lesion burden was significantly decreased (*P* < 0.0001) relative to that of DMSO treated mice; whereas, similar lesion burdens were found in Alda-1 and DMSO treated ALDH2*2 mice (Fig. 6H). Combined, these findings suggest that in established endometriosis, Alda-1 pain alleviation is associated with a decreased lesion number and burden in wild type mice and a reduced lesion number, in ALDH2*2 mice.

## Discussion

This study investigated the role of reactive aldehyde detoxification by ALDH2 in endometriosis and its associated pain. In women with endometriosis, elevated peritoneal fluid reactive aldehyde levels are implicated and through accumulation, reactive aldehydes can form tissue protein-adducts and also generate pain (13-19). Our results show that in the proliferative phase, women with severe (IV) peritoneal endometriosis, have decreased ALDH2 activity and expression in the endometrium (eutopic and ectopic) compared to endometrium (eutopic) of women without endometriosis, suggesting altered ALDH2 activity may underly endometriosis disease pathology. Further supporting this hypothesis, in women with endometriosis, 4-HNE adduct formation was increased in the endometrium (eutopic and ectopic) of women with endometriosis compared to the endometrium (eutopic) of women without endometriosis, suggesting protein adducts may form as a result of the decreased reactive aldehyde detoxification by ALDH2.

Using a rodent model of endometriosis and an ALDH2*2 knock-in mouse, with reduced ALDH2 activity and expression and similar basal levels of 4-HNE adducts, relative to wild type mice, we provide evidence that early in endometriosis development, decreased ALDH2 activity accelerates lesion development and exacerbates pain-associated behavior. We further determined that increasing ALDH2 activity with the enzyme activator, Alda-1, could prevent early lesion development. Once endometriosis was established, we determined that increasing ALDH2 activity with Alda-1 treatment, could alleviate endometriosis pain-associated behavior which was associated with a decreased lesion number in both wild type and ALDH2*2 mice and a decreased lesion burden number in wild type mice. Combined, our findings from women with endometriosis and a preclinical rodent model of endometriosis, support our overall hypothesis that aberrant reactive aldehyde detoxification by ALDH2, underlies endometriosis and its associated pain.

Overall, our findings suggest that targeting ALDH2 enzyme activity may be effective for the alleviation of endometriosis-associated pain, particularly primary/local abdominal pain. Early on, ALDH2*2 endometriosis mice, with reduced ALDH2 activity, develop larger lesions and exacerbated abdominal pain-associated behaviors, relative to wild type mice, suggesting the lesion’s reduced ability to detoxify reactive aldehydes within the peritoneal cavity, contributes to endometriosis pain-associated behaviors. The decreased ALDH2 activity in the ectopic endometrium (lesions) of the women with peritoneal endometriosis, further supports this. With reduced ALDH2 activity and therefore, reduced reactive aldehyde detoxification, elevated levels of reactive levels such as 4-HNE can influence lesion innervation, to induce pain.

In women and rodent models of endometriosis, lesions develop a sensory and sympathetic nerve supply which opens a 2-way line of communication between the lesions and the spinal cord allowing engagement of the nervous system and the generation of pain (21-26). Often co-expressed on sensory nerves, are the nociceptive ion channels transient receptor potential ankyrin 1 (TRPA1) and transient receptor potential vanilloid 1 (TRPV1), where they integrate noxious stimuli and generate pain in inflammatory conditions (27, 28). In women with endometriosis, lesion TRPV1 and TRPA1 mRNA and immunoreactivity are upregulated and positively correlated with painful symptom severity, compared to the eutopic endometrium of controls with no pain (30). Moreover, mRNAs encoding TRPA1 and TRPV1 are increased in the peritoneum of women with endometriosis and chronic pelvic pain, compared to the peritoneum of healthy women with no pain (31). Therefore, elevated reactive aldehydes, such as the endogenous TRPA1 agonist 4-HNE, implicated in the peritoneal fluid of women with endometriosis, could activate TRPA1 channels on sensory nerve fibers in lesions and/or the peritoneal cavity, in concert with TRPV1 channels, to drive increased pain signaling (31). If true, increasing reactive aldehyde detoxification by ALDH2 with Alda-1, may reduce endogenous reactive aldehyde sensitization of TRPA1, to reduce pain. Further, topical or local administration (i.p.) of Alda-1 may alleviate endometriosis-associated primary abdominal/pelvic pain, reducing the risk of side effects common in oral/systemic routes of drug administration.

Our findings further suggest that ALDH2 activity influences lesion development and maintenance which has implications for surgery-based treatment strategies for endometriosis-associated pain. In mice, decreased ALDH2 activity exacerbated lesion development whereas increasing ALDH2 prevented lesion development. Once established, increasing ALDH2 activity decreased endometriosis lesion number and burden. In women with endometriosis, laparoscopic excision of the lesions alleviates pain in ∼50% of cases (32). However, in the ∼50% of women in which surgery is successful, ∼25% have a return of the pain that is frequently accompanied by a return of the lesions (32-34). If our findings are translational in women in which surgery is successful, Alda-1 treatment during and/or after laparoscopic lesion excision may be effective in preventing the return of the lesions and/or pain. In women in which surgery is unsuccessful in pain alleviation or women that chose not to have surgery, Alda-1 treatment could reduce lesion number and/or total lesion burden to reduce pain.

Findings from this study also provide potential insight into mechanisms underlying endometriosis etiology. The decreased ALDH2 activity and expression in the eutopic and ectopic endometrium of women with endometriosis, suggests that ALDH2 activity and expression differences may help explain why 90% of women have retrograde menstruation but only 10% of women develop endometriosis. The decreased reactive aldehyde detoxification by ALDH2 in women with endometriosis is likely key in reactive aldehyde accumulation, promoting an environment of oxidative stress and therefore, extrauteral endometrium (lesion) implantation and survival of (10, 11). The concomitant increase in 4-HNE adducts in the endometrium of women with endometriosis, provides preliminary support for 4-HNE adduct formation as a diagnostic biomarker, in the proliferative phase. The gold standard for endometriosis diagnosis is laparoscopic lesion visualization preferably with histological confirmation, which is invasive and expensive (9). Therefore, biopsy of the eutopic endometrium or endometrial curettage with subsequent 4-HNE analysis, may serve as a less invasive diagnostic biomarker. Overall, a better understanding of the differences between women with and without endometriosis are critical in the development of more effective treatment and diagnostic strategies.

Limitations of our study include human tissue sample size. Due to the limited samples specific to our request, only 5 (eutopic) samples form women without endometriosis and 10 samples from women with endometriosis (5 ectopic, 5 matched eutopic) were analyzed. Limiting the applicability of our findings, all endometriosis biopsies were from women with peritoneal endometriosis, excluding other endometriosis subtypes (ovarian endometriomas, deep infiltrating endometriosis) involving different etiologies that may require different diagnostic and treatment strategies (35). Our biopsies were all from the proliferative phase of the menstrual cycle to control for differences within the menstrual cycle; however, we did not control for potential gene expression differences within the days of the proliferative phase (36). In a larger future study, we will analyze additional endometriosis subtypes and days within the proliferative phase. Our analysis will also include biopsies from additional races, particularly East Asians of which ∼50% have the ALDH2*2 genetic variant with reduced ALDH2 activity that our ALDH2*2 knock in mouse mimics. Our findings in ALDH2*2 mice may be translatable to women with the ALDH2*2 variant; however, a direct link between the E487K mutation and endometriosis must be established in future studies. Further, cell specific ALDH2 characterization in endometrial tissue will performed, as this is yet to be explored.

Animal study limitations include the inability to distinguish between the contribution of reduced uterus and liver ALDH2 activity to endometriosis pathology, which would require a conditional uterine knockout model. Although desirable, it was beyond the scope of the original study to determine the relationship between ALDH2 and 4-HNE adduct formation in the endometriosis mouse model. Our primary objective was to establish aberrant ALDH2 activity in women with endometriosis and then, in a preclinical endometriosis model, manipulate ALDH2 activity to determine its role in lesion development and pain-associated behaviors. However, other studies have established the relationship between ALDH2 activity, 4-HNE, and Alda-1 in other tissues (37). Future studies will further test our hypothesis and assess the relationship between ALDH2, 4-HNE, and Alda-1 in women and mice with endometriosis.

Overall, our findings establish the ALDH2 enzyme as a novel target for endometriosis and its associated pain and Alda-1, as a potentially therapeutic. Further, we provide preliminary support for the development of increased 4-HNE protein adduct formation, in the endometrium, as a biomarker to reduce diagnostic delay and years of suffering in women with endometriosis.

## Materials and Methods

### Study design

The primary objective of this study was to evaluate the role of ALDH2 reactive aldehyde detoxification in endometriosis. First, ALDH2 activity, ALDH2 expression, and 4-HNE adduct formation were assessed in the endometrium of women with endometriosis (lesion and eutopic) compared to women without endometriosis (eutopic) to determine how ALDH2 activity was regulated. We then biochemically characterized an ALDH2*2 knock-in mouse and used a preclinical endometriosis model to determine if ALDH2 activity influences disease development. Subsequently, we incorporated the ALDH2 activator Alda-1 and behavioral assessments to determine if increasing ALDH2 activity could prevent endometriosis development and/or alleviate pain-associated behaviors. For the animal studies, the number per group was based on previous experience with the disease model and a power analysis. Mice were randomly placed in control and treatment groups and experimenters were blinded to conditions. All sample processing in this study was performed concurrently on experimental and control groups using identical methods. Experimenters quantifying results were blinded. Statistical tests and number of animals or replicates for each experiment are included in the figure legends.

### Human tissue samples

In total, fifteen proliferative phase endometrial tissue samples total were obtained from women with histologically confirmed, severe (stage IV) peritoneal endometriosis at laparoscopy (n=5 eutopic, n=5 patient matched ectopic peritoneal lesions) and from women found to be free of endometriosis at surgery (n=5 eutopic). All subjects were Caucasian, non-smokers (except one past smoker), had regular menstrual cycles, and had not received steroid hormone medications within 3 months of endometrial sampling (Table 1). The mean age of participants was 37.8 ± 3.44 years for the endometriosis samples and 41.4 ± 2.25 years for the non-endometriosis samples. Women without endometriosis at surgery were undergoing hysterectomy or gynecological surgery for a benign condition. All endometrial samples were obtained from the University of California-San Francisco (UCSF) National Institute of Health (NIH) Human Endometrial Tissue and DNA Bank. All samples procured by the Tissue Bank are obtained after written informed consent under an approved protocol by the UCSF Committee on Human Research. UCSF sample acquisition and storage are by established standard operating procedures (SOPs) (38). Menstrual cycle phase was assigned as proliferative phase endometrium (PE) by endometrial histology according to the Noyes criteria (39). Severe endometriosis (stage IV disease) was defined in accordance with the Revised American Fertility Society (rAFS) classification system in which disease stage is graded on a scale of I (minimal), II (mild), III (moderate), to IV (severe) based on endometrial tissue location, amount, depth, and associated adhesions (40). Peritoneal endometriosis was defined as biopsy-proven serosal implant.

### ALDH2*2/*2 knock-in mouse and vaginal cytology

Animal subjects were virgin female ALDH2*2 knock-in (n=107) and C57BL/6 wild type (n=107) litter mate mice, 6-8 weeks. Mutant mice were homozygous ALDH2*2 (ALDH2*2/*2) knock-in mice on a C57BL/6 background generated by replacing the mouse wild type ALDH2 allele with a mouse E487K mutant ALDH2 allele by homologous recombination (Fig. 2). Compared to wild type mice, the ALDH2*2 knock-in mouse has a single amino acid substitution in which adenine is substituted for guanine at the first base pair of codon 487. As a result, there is an amino acid change from glutamic acid (Glu, E) to lysine (Lys, K) that is equivalent to the E487K substitution in the ALDH2*2 human variant, in which ALDH2 enzyme activity is significantly decreased compared to that of wild type mice (41). Founder mice were back-crossed to the C57BL/6 background for at least 7 generations to achieve a homogeneous genetic background as previously described (41). All mice were group housed in a temperature-controlled room (22°C), in Innovive cages lined with chip bedding and *ad libitum* access to rodent chow and water. Housing was in environmentally controlled conditions (room temperature ∼22°C; 12-h light/dark cycle, with lights on at 07:00). Reproductive status was determined by vaginal lavage performed ∼2h after lights on using traditional nomenclature for the four estrous stages of proestrus, estrus, metestrus, and diestrus (42). To control for the potential confound of vaginal lavage as an acute stressor, on behavioral assessments, animals were handled daily. The study and procedures were approved by the Animal Care and Use Committee as Stanford Univeristy protocol #32871 and Emory University protocol #201900201. All laboratory animal experimentation adhered to the National Institute of Health (NIH) Guide for the Care and Use of Laboratory Animals.

### Endometriosis surgery (ENDO)

At ∼10 weeks of age, in the estrous stage of diestrus, mice were induced with endometriosis (ENDO) based on the rat model protocol originally developed by Vernon and Wilson and slightly modified by Cummings and Metcalf (43, 44). Briefly, mice were anesthetized with isoflurane (1-3%) and placed on a heating pad to maintain body temperature (37C). An off-midline (left side) incision was made through the skin and muscle layer to expose the pelvic and abdominal organs. A ∼1-cm segment of mid-left uterine horn with its attached fat was ligated proximally and distally with suture. Then, the uterine horn was excised, and the associated fat removed. Three, 2 mm x 2 mm pieces of the excised uterus (minus the fat) were sewn onto alternate mesenteric arteries that supply the caudal small intestine starting from the caecum using 4.0 nylon sutures. After it was confirmed that no bleeding was occurring in the abdominal cavity, the muscle layer was closed with chromic gut suture, skin incision closed with silk suture, and mice closely monitored during recovery.

### Osmotic pump implantation and drug delivery

For Alzet osmotic pump implantation, an incision was made at the nape of the mouse neck and the pump implanted immediately below the skin layer and closed with silk suture for subcutaneous continuous delivery of Alda-1 (5mg/kg/day), or as a control solvent only (50% polyethylene glycol-PEG and 50% dimethyl sulfoxide-DMSO by volume). Pumps were filled and then primed in 0.9% sterile saline at 37°C for ∼24 hours prior to implantation.

### Behavioral assessments

Behavior tests included measures of abdominal licking, mechanical nociception, thermal nociception, locomotor activity, and exploratory behavior, all previously shown to be altered in association with endometriosis pain-associated behaviors (45-48). Abdominal licking has been used an indicator of abdominal pain in various viscero-specific pain models (46, 49-51). Therefore, the number of times the mouse licked, groomed, barbered the abdominal region was recorded as an indicator of local abdominal discomfort and indicator of primary/local pain. This test was done in a modified home cage open field setting with bedding and the mouse first allowed a minimum of 10 minutes to acclimate. To measure changes in nociception, hind paw withdrawal threshold and thermal latency were assessed as a secondary or referred pain-associated behavior. To assess mechanical nociception, von Frey fibers were used ranging in force from 0.004 to 5.49g using the Dixon modified up and down method (52, 53). Mice were placed individually in a plexi-glass chamber on an elevated mesh screen stand and allowed to acclimate for a minimum of 10 minutes. Von Frey hairs were applied perpendicularly to the mouse hind paw plantar surface until the hair bowed and then held for approximately 3 seconds. The mechanical threshold required to elicit a paw withdrawal (50% paw withdrawal threshold) was determined. To assess for changes in thermal nociception, Hargreaves method was used to determine latency to paw withdrawal from a focused heat light source using a commercial Plantar Test Analgesia Meter (IITC Life Science) (54). Mice were placed individually in plexi-glass chambers on a glass platform and allowed to acclimate for a minimum of 10 minutes. The heat stimulus was delivered to the plantar region of the mouse hind paw with an active intensity of 30%. Reaction time was measured in .01 second increments with a cut off time of 10 seconds. A minimum of 30 seconds separated each hind paw test.

To assess for changes in locomotor activity, mice were individually placed in an automated Opto-Varimex activity monitor (Columbus Instruments) with optical beam sensors and total (ambulatory and non-ambulatory), ambulatory (does not include stereotypic non-ambulatory behavior e.g. grooming, digging), and vertical (rearing) counts were recorded. To assess for changes in exploratory behavior, a cardboard tunnel was included in the modified home cage open field setting. The number of cardboard tunnel entries, amount of time spent in the tunnel (s), and number of times the mouse climbed on top of the tunnel was recorded.

Abdominal licking, locomotor activity, and exploratory behavior were assessed in separate 5 minutes sessions. Behavioral tests were run in the order thought to be least stressful test to most invasive [home cage assessments (abdominal licking, exploratory behavior), locomotor activity, Von Frey, and Hargreaves] as test order may influence behavior (55). Two experimenters, SM and MV, performed behavioral assessments (inter-observer reliability, r > 0.90) blinded to treatment. Because male experimenters can produce pain inhibition, both experimenters were females (56). Because endometriosis is an estrogen-dependent condition and pain-associated behaviors are influenced by estrogen, all behavioral data is reported in the estrous stage of proestrus (42, 57-59). Behavioral parameters were assessed twice in proestrus twice during each 2-week time period applicable (baseline, post-endometriosis, and post-treatment). Testing days were ∼day 5 and day 13 of each 2-week time period.

### Mouse tissue collection

At the time of sacrifice under isoflurane anesthesia, the abdominal cavity was opened and examined. When applicable the area where the eutopic uterus auto-transplants were previously sewn was investigated and sutures located to identify and measure the lesions *in situ*. The ectopic lesions, eutopic uterus, and liver (control) were harvested, immediately frozen in dry ice, and stored at −80C. After tissue harvesting, mice were sacrificed.

### Mouse ectopic lesion measurement

To assess for differences in lesion size, total lesion burden was first determined. To do this, the largest and smallest diameter of each lesion was multiplied to give a value (most lesions have an ovoid shape) and then values from each lesion added to obtain a total number, the total lesion burden (24, 60). The total lesion burden was divided by the number of lesions formed to give the average lesion area for comparison between groups.

### Tissue processing and analysis

Mouse and human tissue samples were homogenized in sucrose mannitol buffer, pH 7.4 (210 mM mannitol, 70 mM sucrose, 1.0 mM EDTA, 5.0 mM MOPS) with protease and phosphatase inhibitors. Briefly, tissue homogenates were centrifuged to remove cellular debris and the supernatant was retained as the whole cell fraction for the analysis. Samples were immediately stored at −80C until further analysis.

### Western blot analysis

Total protein concentration was determined by the Bradford assay according to the manufacturer’s protocol. Equal amounts of protein (30 µg) for each sample were separated by SDS-PAGE on 4-15% polyacrylamide gels and transferred to PVDF Membranes (Bio-Rad) and probed overnight at 4 °C for specific antibodies against ALDH2 (Santa Cruz Biotechnology), 4HNE (Alpha diagnostics), and actin (Cell Signaling). The next day, membranes were washed and incubated a with horseradish-peroxidase (HRP) linked secondary antibody (anti-goat, Santa Cruz Biotechnology or anti-rabbit, Cell Signaling) for 1 hour at room temperature. Membranes were again washed and bound antibody was detected by enhanced chemiluminescence. Images were acquired by using an Azure Biosystems c300. Image-J (NIH) software was used for the relative expression and densitometry analysis. Relative protein expressions were normalized to actin (loading control).

### Enzymatic activity assay

Cofactor and substrate (NAD^+^ and 25mM acetaldehyde) were added to the reaction buffer containing homogenate tissue and the conversion of NAD+ to NADPH over time monitored using a spectrophotometer. For a 1 mL assay, 500 μl of 200 mM NaPPi at final concentration of 200 mM NaPPi in water (pH 9.0 (M.W. 446)), 250 μl of 10 mM NAD^+^ (2.5 mM NAD^+^), 100 μg protein from tissue, and homogenization buffer (0.1 M tris HCL pH 8.0 and 1% triton-X) were added and mixed to make 1 mL total volume. Absorbance (O.D.) was measured at A340 nm for 3 min. Then, 2.5 μl of 10 mM acetaldehyde (f.c., 25 mM) was added to the cuvette and the absorbance measured for an additional 15 minutes. ALDH2 activity was converted to µmole NADH/min/mg of protein. As a blank control, cuvettes without tissue/sample or acetaldehyde were used. Data presented is absorbance measured during the first *2* minutes after acetaldehyde substrate was added.

### Statistical analysis

To achieve at least a 20% minimal difference between groups for a power of 95% with α<0.05 and β<20% a minimum of 6 mice/group were used. Data are expressed as mean ± SEM. For data with only two groups, a two-tailed Student’s t-test was used. For data containing more than two groups, a one-way or two-way analysis of variance (ANOVA) was used, followed by Tukey post hoc test as appropriate. Statistical meaningful differences were assumed for p<0.05. All statistical analysis was performed using GraphPad 8.12.

## Acknowledgments

Tissue samples were provided by the NIH P50 National Centers in Translational Research in Reproduction (NCTRI) Human Endometrial Tissue Bank and DNA Bank at UCSF, funded under NIH HD055764-11(LCG). The authors thank Drs. Linda Giudice, Rona Giffard, Eric Gross, and Laure Aurelian for helpful discussions during the study. The authors also thank Drs. Julie Christianson and Neil Sidell for helpful input during the writing of the manuscript and Daria Mochly-Rosen for ALDH2*2 knock-in mouse access.

## Author Contributions

SLM conceived the presented idea, performed molecular and behavioral experiments and rodent surgeries, analyzed the data, made the figures, and wrote the manuscript with input from all authors. PSR optimized molecular protocols, performed molecular experiments, and analyzed data. MV performed molecular and behavioral experiments and made illustrations. All authors read and approved the final manuscript.

## Data and materials availability

All data from this study is provided in the manuscript or supplemental material. Materials are available upon reasonable request.

---

**Fig S1.**
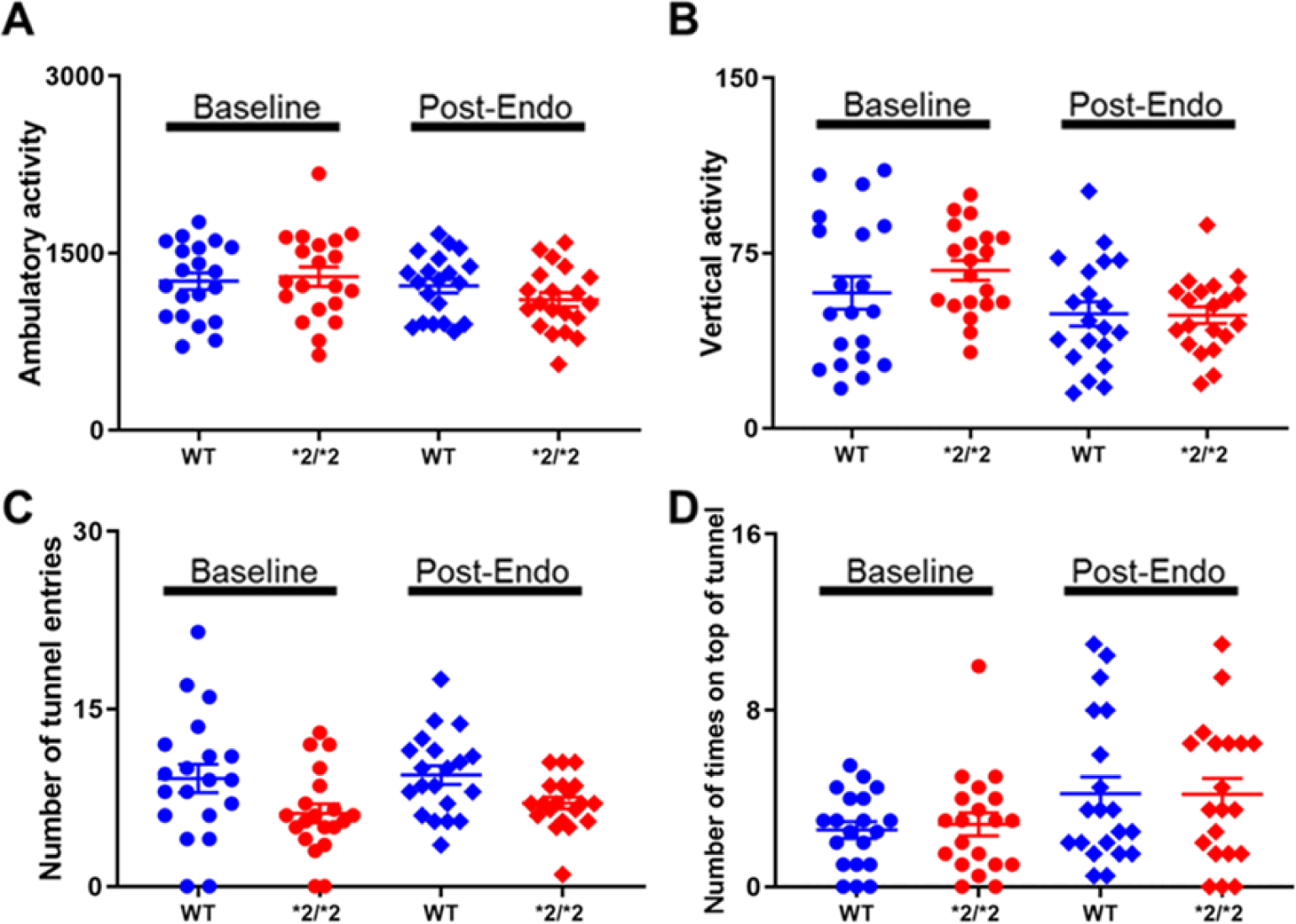
Effects of decreased ALDH2 activity on locomotor activity and exploratory behavior in endometriosis mice. Behavioral assessments were made post-endometriosis (n=2) and compared to baseline (n=2) in wild type (n=20) and ALDH2*2 mice (n=20) in proestrus. Locomotor activity and exploratory behavior were assessed in 5 minute sessions and included ambulatory activity (A); vertical activity (B); number of tunnel entries (C) and number of times the mouse climbed on top of the tunnel (D). All data are expressed as mean ±SEM. Assessed by two-way ANOVA with Tukey’s post hoc test, blue and red data points indicate wild type mice, red data points indicate ALDH2*2/*2 mice, circles baseline assessments, diamonds post-endometriosis assessments.

**Fig S2.**
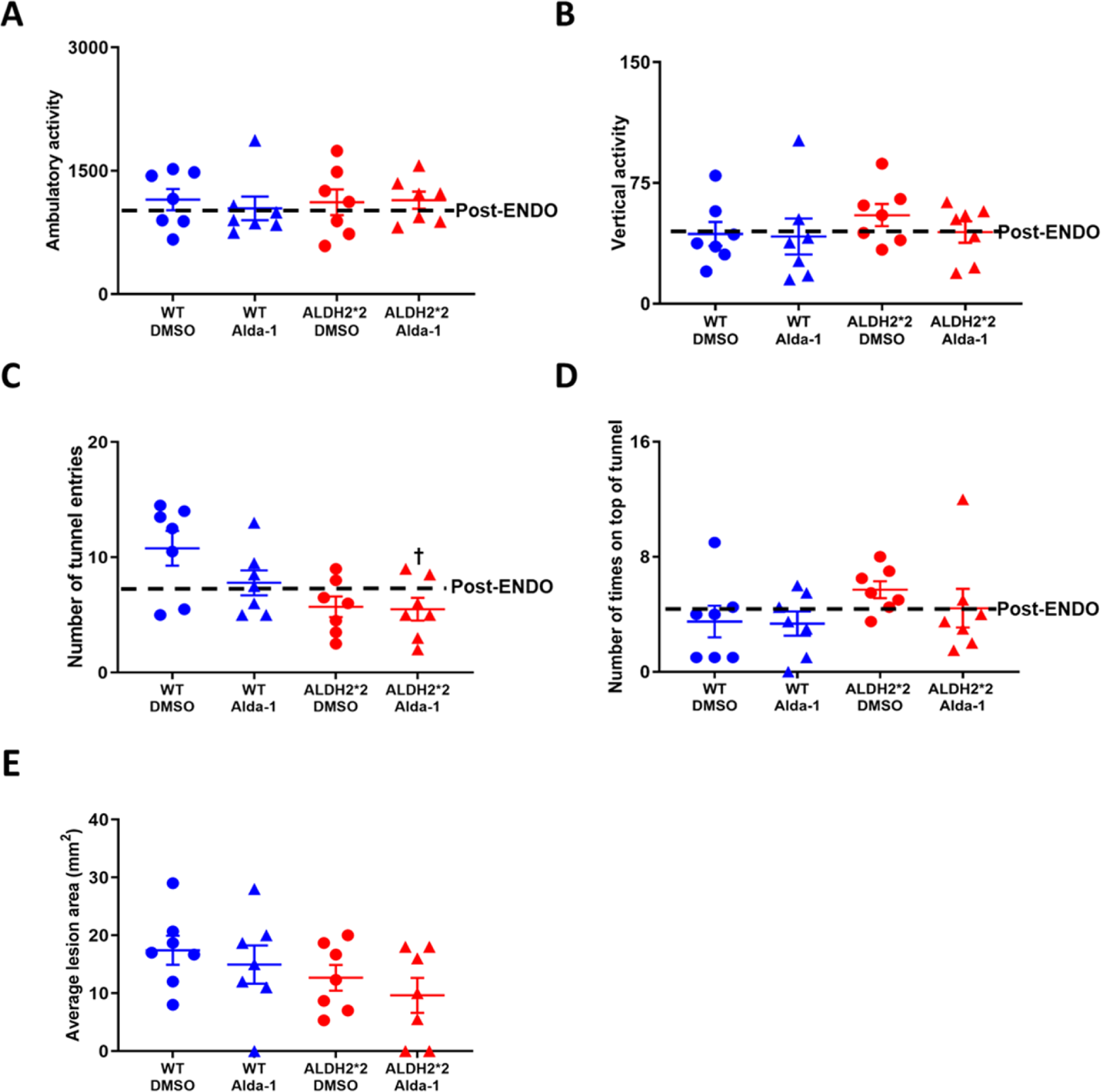
Effects of increased ALDH2 activity on locomotor activity and exploratory activity in endometriosis mice. Endometriosis surgery performed in wild type (n=14) and ALDH2*2 mice (n=14). At day 14 post-endometriosis, Alda-1 treatment (5mg/kg/day or 50-50 DMSO-PEG control) began and continued for 2 weeks via Alzet osmotic pump (n=7/group/genotype). In proestrus, behavioral parameters were assessed and post-treatment (n=2) assessments compared to post-endometriosis (n=2). Locomotor activity and exploratory behavior were assessed in 5 minute sessions and included ambulatory activity (A) and vertical activity (B), number of tunnel entries (C) and the number of times the mouse climbed on top of the tunnel (D). Mice were sacrificed and average lesion area (A) compared between groups. All data are expressed as mean ±SEM. Behavioral data assessed by two-way ANOVA, followed by Tukey post hoc test. Lesion data assessed by one-way ANOVA, followed by Tukey post hoc test. Blue data points indicate wild type mice, red data points indicate ALDH2*2 mice, circles DMSO treated, and triangles Alda-1 treated. *p<0.05 *vs*. wild type baseline

